# *Clostridioides difficile* SpoVAD and SpoVAE interact and are required for DPA packaging into spores

**DOI:** 10.1101/2021.07.28.454260

**Authors:** Marko Baloh, Joseph A. Sorg

## Abstract

*Clostridioides difficile* spores, like the spores from most endospore-forming organisms, are a metabolically dormant stage of development with a complex structure that conveys considerable resistance to environmental conditions, *e.g.*, dry heat. This resistance is due to the large amount of dipicolinic acid (DPA) that is packaged into the spore core, thereby replacing the majority of water. DPA is synthesized by the mother cell and its packaging into the spore core is regulated by the *spoVA* operon that has a variable number of genes, depending on the organism. *C. difficile* encodes 3 *spoVA* orthologues, *spoVAC, spoVAD,* and *spoVAE.* Prior work has shown that *C. difficile* SpoVAC is a mechanosensing protein responsible for DPA release from the spore core upon the initiation of germination. However, the roles of SpoVAD and SpoVAE remain unclear in *C. difficile.* In this study we analyzed the roles of SpoVAD and SpoVAE and found that they are essential for DPA packaging into the spore, similar to SpoVAC. Using split luciferase protein interaction assays we found that these proteins interact, and we propose a model where SpoVAC / SpoVAD / SpoVAE proteins interact at or near the inner spore membrane, and each member of the complex is essential for DPA packaging into the spore core.

**Importance:** *C. difficile* spore heat resistance provides an avenue for it to survive the disinfection protocols in hospital and community settings. The spore heat resistance is mainly the consequence of the high DPA content within the spore core. By elucidating the mechanism by which DPA is packaged into the spore core, this study may provide insight in how to disrupt the spore heat resistance with the aim of making the current disinfection protocols more efficient at preventing the spread of *C. difficile* in the environment.

## Introduction

*Clostridioides difficile* is a Gram-positive, spore-forming, strictly anaerobic bacterium that is opportunistically pathogenic in humans. According to the most recent report by the Centers for Disease Control and Prevention, issued in 2019, it is estimated that 223,900 cases of *C. difficile* occurred in 2017 (1, 2). Of these, 12,800 deaths can be directly attributed to *C. difficile*, with the majority of deaths occurring among people aged 65 and older. In recent years, there has been an emergence of antibiotic-resistant strains, as well as strains with increased virulence making *C. difficile* a leading cause of hospital-associated infections with an estimated $5 billion in annual treatment-associated cost for *C. difficile* infections (CDI) in US hospitals alone (3, 4).

*C. difficile* infections are initiated upon disruption to the normally protective microbiota, commonly due to broad spectrum antibiotic use (5–8). Antibiotics are prescribed to treat CDI (*i.e.* vancomycin or fidaxomicin) but patients frequently relapse with recurring CDI due to the continued disruption to the colonic microbiome and the presence of antibiotic-resistant spores that remain in the gastrointestinal tract or in the surrounding environment (9). Though the *C. difficile* vegetative form is the disease-causing agent, it is the spore form that is the infective agent due to its ability to survive outside of the host in the aerobic environment. *C. difficile* spores are structurally complex, composed of several distinct layers that are broadly similar to spores produced by other endospore-forming organisms, *e.g., Bacillus subtilis* (10, 11). In the spore core the majority of the water is replaced by pyridine-2,6-dicarboxylic acid (dipicolinic acid [DPA]), chelated with calcium (Ca-DPA), which provides extreme heat resistant properties (12, 13). The core is surrounded by an inner membrane that has low permeability, preventing water, and potentially DNA-damaging molecules, from entering the core (14, 15). Surrounding the inner spore membrane is a germ cell wall that will become the cell wall of the vegetative cell, post-germination. Surrounding the germ cell wall is a thick layer of cortex peptidoglycan. In the cortex, a proportion of muramic acid residues is converted to muramic-δ-lactam (16). The muramic-δ-lactam residues are the targets of cortex lytic enzymes during spore germination (17, 18). The cortex is surrounded by the outer spore membrane and by the coat, which provides the spore with protection from environmental insults and decontaminants (19). In some endospore-forming bacteria, the outermost layer of spores is an exosporium layer, which may have a role in adherence to host intestinal epithelial cells and other surfaces (20, 21).

Spores are metabolically dormant until the detection of germinants by germinant receptors. Germinant recognition initiates a cascade of events that irreversibly commits the spore to the germination pathway (22–24). This event is followed by the release of DPA from the core and degradation of the cortex by spore cortex lytic enzymes (SCLE), either simultaneously or sequentially, depending on the organism (18, 25, 26). In the model spore-forming bacterium, *B. subtilis*, SCLEs can be activated by exogenously added Ca-DPA or the release of Ca-DPA from the core, meaning that DPA release precedes cortex hydrolysis (18). However, in *C. difficile* these two early germination steps are inverted and cortex hydrolysis precedes DPA release (12, 27).

In *C. difficile,* activation of the SleC cortex lytic enzyme leads to the release of DPA stores from the spore core in exchange for water. In *B. subtilis* the proteins encoded by the *spoVA* operon (SpoVAA-AB-AC-AD-AEa-AEb-AF) play a role in DPA packaging and release, but *C. difficile* encodes only 3 orthologues: *spoVAC, spoVAD*, and *spoVAE*. *B. subtilis* SpoVAC is a mechanosensing protein, similar to its role in *C. difficile* (27, 28). In *B. subtilis,* SpoVAD binds to DPA and is likely important in packaging of DPA into the developing spore and a *spoVAEa* mutation causes only a slight germination defect, while the role of *spoVAEb* is unclear (29, 30). Herein, we show that *C. difficile spoVAD* and *spoVAE* are essential for DPA packaging into the spore. Because in *B. subtilis* the proteins of the *spoVA* operon are hypothesized to form a membrane channel enabling the uptake of DPA into the forming spore during sporulation and release of DPA from the spore during germination, we speculated that this also may be the case in *C. difficile* (30, 31). Using a split-luciferase reporter system we find that indeed some of the products of the *spoVA* operon interact. This led us to propose a model of a SpoVAC / SpoVAD / SpoVAE interaction that describes their roles in DPA packaging and release during *C. difficile* spore formation and germination, respectively.

## Results

### *spoVAD* and *spoVAE* are required for packaging of DPA into the spore

Prior research on *spoVAC* found that a *spoVAC* deletion resulted in spores that contain only 1% of the wild-type level of DPA (27, 32). To understand the contribution of *spoVAD* and *spoVAE* to sporulation and DPA packaging, we used an established p*yrE*-based allelic exchange system (33) to create *spoVAD* or *spoVAE* deletion strains in the *C. difficile* CRG2359 strain, restored the *pyrE* deletion, and then analyzed for their DPA content using terbium fluorescence (34). Spores derived from *C. difficile spoVAD* and *spoVAE* mutant strains contained approximately 1% of the wild-type levels of DPA (Figure 1A). This amount of DPA content could be restored to wild type levels by expressing *spoVAD* or *spoVAE*, *in trans,* from a plasmid. These results suggest that *spoVAD* and *spoVAE*, like *spoVAC*, are required for DPA packaging into the *C. difficile* spore, similar to their roles in *B. subtilis* (29, 30).

**Figure 1.**
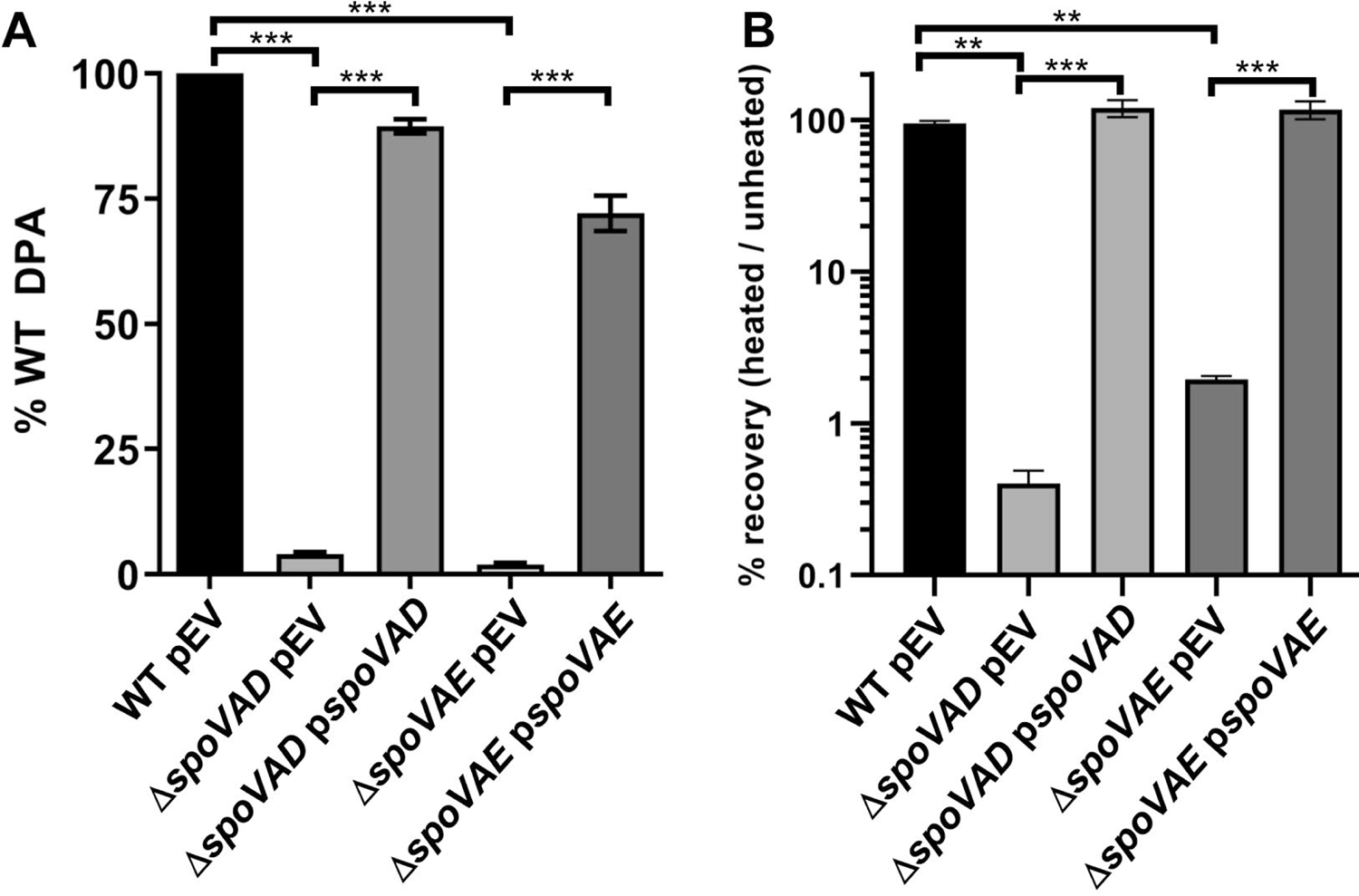
*spoVAD* and s*poVAE* are required for DPA packaging into the spore. (A) Equal amounts of spores purified from *C. difficile* strains RS19 (WT), MB03 (Δ*spoVAD*), and MB04 (Δ*spoVAE*) were boiled for 20 minutes and DPA amounts quantified by Tb^3+^ fluorescence. Values are reported as percentage of the WT DPA content. WT and the mutants were transformed with an empty vector (pEV; pJS116) to account for the effects of the presence of the plasmid. (B) 1×10^8^ spores were serially diluted and plated on BHIS plates supplemented with TA, before and after heating at 65 °C for 30 minutes. The values are reported as percentage of heated spores that formed colonies, compared to unheated spores. The data represents results from 3 independent assays and the error bars represents the standard error of the mean. ** indicates p<0.001, *** indicates p<0.0001 as determined by one-way ANOVA using Tukey’s multiple comparisons test.

Next, we tested if the absence of DPA led to a loss of heat resistance to spores derived from the mutant strains. When spores derived from the *C. difficile* Δ*spoVAD* and Δ*spoVAE* strains were heated at 65 °C for 30 minutes and plated on BHIS medium supplemented with taurocholic acid (a potent spore germinant), we observed a >90% decrease in the number of colony forming units compared to the unheated samples or to the samples containing complementation plasmids (Figure 1B). These results show that spores derived from the *C. difficile* Δ*spoVAD* and Δ*spoVAE* mutant strains are heat sensitive due to the lack of DPA or that heat treatment blocks an early germination event.

### *spoVAE* may contribute to *C. difficile* spore germination

*C. difficile* spore germination is activated upon binding of certain bile acid and certain amino acid germinants to receptors (23, 34–37). This results in the irreversible initiation of germination and the release of the majority of DPA from the spore core. Another indicator of germination is the change of optical density of the spore solution. This assay takes advantage of the transition of dormant spores from a phase-bright, dormant, state to a phase-dark, germinated, state. In both assays, equal numbers of spores derived from *C. difficile* RS19 [*C. difficile* CRG2359 with a restored *pyrE* (36)], *C. difficile* Δ*spoVAD*, or *C. difficile* Δ*spoVAE* strains containing empty vectors or the complementation constructs, were suspended in buffer alone or buffer supplemented with the germinants taurocholate and glycine. Subsequently, the release of DPA and the change in OD_600_ values were measured over a period of 1 hour.

Due to the reduced abundance of DPA packaged into the spores, *C. difficile* Δ*spoVAD* spores release very little DPA (Figure 2A). This phenotype can be complemented by the expression of the wild type copy of the gene *in trans.* In the optical density-based germination assay we observed only a small, ∼10%, reduction in OD_600_ values for *C. difficile* Δ*spoVAD*-derived spores. The spores derived from the wild type and the complemented strain germinated normally (Figure 2B). Spores derived from the *C. difficile* Δ*spoVAE* strain also released very little DPA, and this phenotype was only partially complemented to wildtype levels (Figure 2C). In the optical density germination assay, *C. difficile* Δ*spoVAE*-derived spores showed an intermediate phenotype, with an ∼30% reduction in OD_600_ values, which can again only be partially complemented to wild-type levels (Figure 2D). Despite the lack of DPA, spores derived from the *C. difficile* Δ*spoVAD* and Δ*spoVAE* strains are capable of initiating germination.

**Figure 2.**
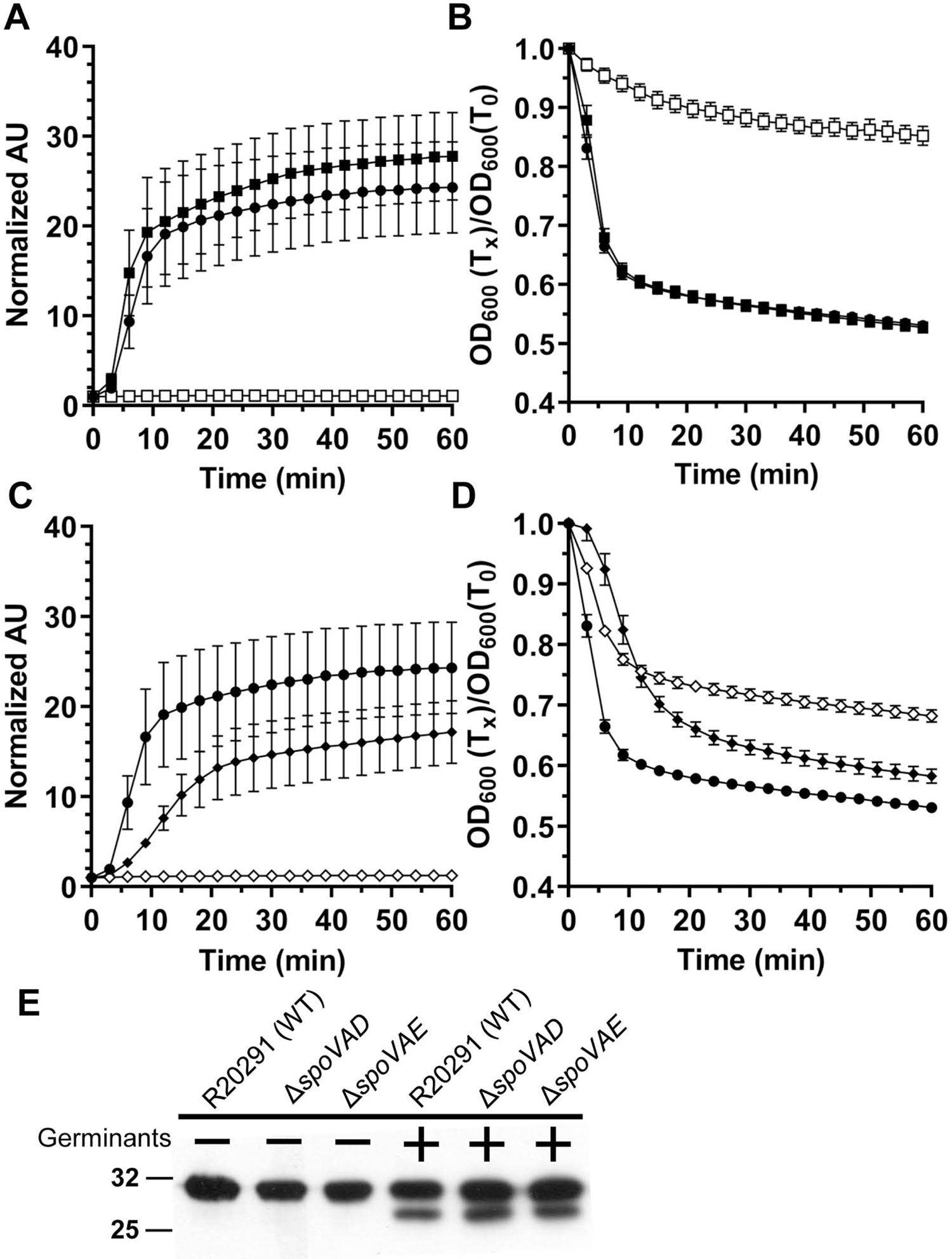
*C. difficile spoVAD* and s*poVAE* mutant spores initiate germination normally. Spores derived from *C. difficile* RS19 pEV (●), MB03 pEV (Δ*spoVAD*; □), MB03 p*spoVAD* (▪), MB04 (Δ*spoVAE*; ◊) and MB04 p*spoVAE* (♦) were purified and germination was quantified by (A and C) DPA release or (B and D) OD_600_. For clarity, every fifth data point is plotted. The data represents results from 3 independent assays and the error bars represent standard error of the mean. (E) 10^8^ spores derived from *C. difficile* CRG2359 (WT), MB03 (Δ*spoVAD*), and MB04 (Δ*spoVAE*) were incubated in buffer alone or in buffer supplemented with the germinants taurocholate and glycine and then boiled in sample buffer, the proteins resolved by SDS-PAGE, and probed with anti-SleC antibody. Inactive pro-SleC corresponds to ∼32 kDa band while activated SleC is ∼29 kDa.

Western blot analysis showed that the cortex lytic enzyme SleC, required for the initiation of *C. difficile* spore germination, was activated to similar levels in the spores derived from the mutant and the wild type strains (Figure 2E). These results suggest that spores derived from *C. difficile* Δ*spoVAD* and Δ*spoVAE* strains activate SleC normally but that the *C. difficile* Δ*spoVAE* mutant strain has a small defect in germination.

### Testing the interaction of the *C. difficile* SpoVAC, SpoVAD, and SpoVAE proteins

Our findings of *C. difficile spoVAD* and s*poVAE* mutant phenotypes indicate that deletion of either gene results in the absence of DPA in the spore core. Prior research has shown a similar phenotype in a *C. difficile spoVAC* mutant (27, 32). In both *B. subtilis* and *C. difficile* the SpoVAC protein is a transmembrane protein, hypothesized to be embedded in the inner spore membrane, and acts in a mechanosensing fashion to release DPA from the spore core during germination (12, 28). While SpoVAD is not predicted to have transmembrane domains, we analyzed the *C. difficile* SpoVAE protein sequence with Constrained Consensus TOPology prediction server (CCTOPS) that predicts protein topology as a consensus of 10 different methods, and the results indicated that SpoVAE has several transmembrane domains, consistent with the predicted *B. subtilis* SpoVAE topology (38). The similar phenotype of single mutants and their predicted topology led us to hypothesize that SpoVAC, SpoVAD, and SpoVAE interact and form a complex at the inner spore membrane. To test the potential interaction of these proteins, we used the luciferase protein interaction assay, developed for use in *C. difficile* (39). Briefly, the codon-optimized luciferase reporter was split into two parts, SmBit and LgBit. Each reporter part was translationally fused to the 3’ end of *C. difficile spoVAC*, *spoVAD*, and *spoVAE*, and each fusion pair was expressed under the control of an anhydrotetracycline (aTc) inducible promoter. The plasmids were introduced into the wildtype *C. difficile* R20291 strain and luciferase activity was measured before induction and 1 hour after induction with aTc. As positive controls, we used the full-length luciferase reporter BitLuc, and the HupA-HupA fusions that were originally used to validate the assay (39). As negative controls, we used single s*poVAC* or *spoVAD* or *spoVAE* fusions (to one reporter fragment) while the second reporter fragment remained unfused. We detected a greater than 2-log_10_ increase in luminescence signal after 1 hour of induction for the positive controls BitLuc and HupA-HupA. Interestingly, we observed an increase in luciferase signal for SpoVAC-SpoVAE, SpoVAD-SpoVAC, and SpoVAD-SpoVAE interaction pairs (Figure 3). Though the signal for other interaction pairs after induction was increased, it did not reach statistical significance. In our negative controls the signal did not increase significantly after induction except, notably, in SpoVAD-SmBit fusion where the signal significantly increased for unknown reasons. Nevertheless, the other negative control, SpoVAD-LgBit does not exhibit a significant luciferase signal increase after induction.

**Figure 3.**
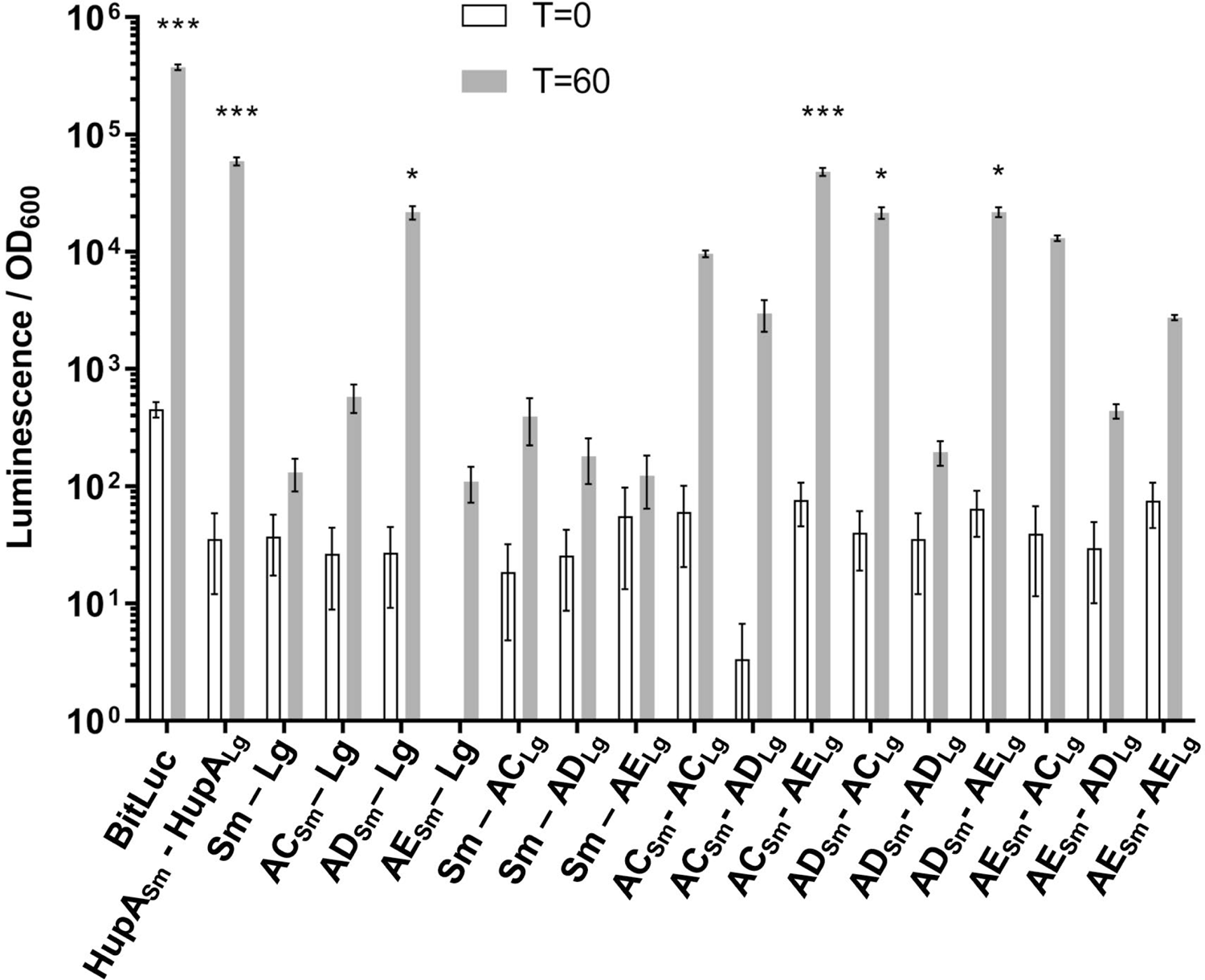
SpoVAC, SpoVAD, and SpoVAE interact *in* vivo *C. difficile* vegetative cells transformed with the indicated plasmids were induced with 200 ng / mL of aTc for 60 minutes. Averages of 3 biological triplicates shown. Optical density-normalized luciferase activity (LU / OD) is shown before induction (white bars) and after 60 minutes (grey bars). Positive interaction was determined by comparison of LU / OD at T = 60 of transformed strains with the negative control (SmBit-LgBit). No significant difference was detected at T=0. * p < 0.05, *** p < 0.0001 as determined by one-way ANOVA using Dunnett’s multiple comparisons test.

Because the packaging of DPA into the spore occurs in the late stages of sporulation (40, 41), we wanted to test for the potential SpoVA protein interactions in cultures that are actively undergoing sporulation. The full length BitLuc control, the negative SmBit-LgBit control, and the interaction partners that showed a significant luciferase signal in induced vegetative cultures (SpoVAC-SpoVAE, SpoVAD-SpoVAC, and SpoVAD-SpoVAE) were cloned into a plasmid and placed under the control of a native *C. difficile spoVAC* promoter (a region of 500 bp upstream of *spoVAC*) and conjugated into the wild-type *C. difficile* R20291 strain. Additionally, we cloned either a SmBit or LgBit luciferase reporter fragment into a segment of the SpoVAC and SpoVAE that is predicted to face the periplasmic space and, therefore, SpoVAD (Supplemental Figure 1S).

The strains were streaked onto sporulation medium and left to grow for 2 days, the period of time that would ensure that the sporulation has commenced, but not fully completed, for the majority of cells in the sample. The cell mass was scraped and resuspended in water, and equal number of cells for each strain were incubated with luciferase substrate and luminescence signal measured (Figure 4). Under these conditions, we observed that SpoVAD and SpoVAE yielded the greatest signal, indicating that during sporulation these two proteins come into close proximity. The other constructs showed a smaller luminescence signal but did not reach statistical significance.

**Figure 4.**
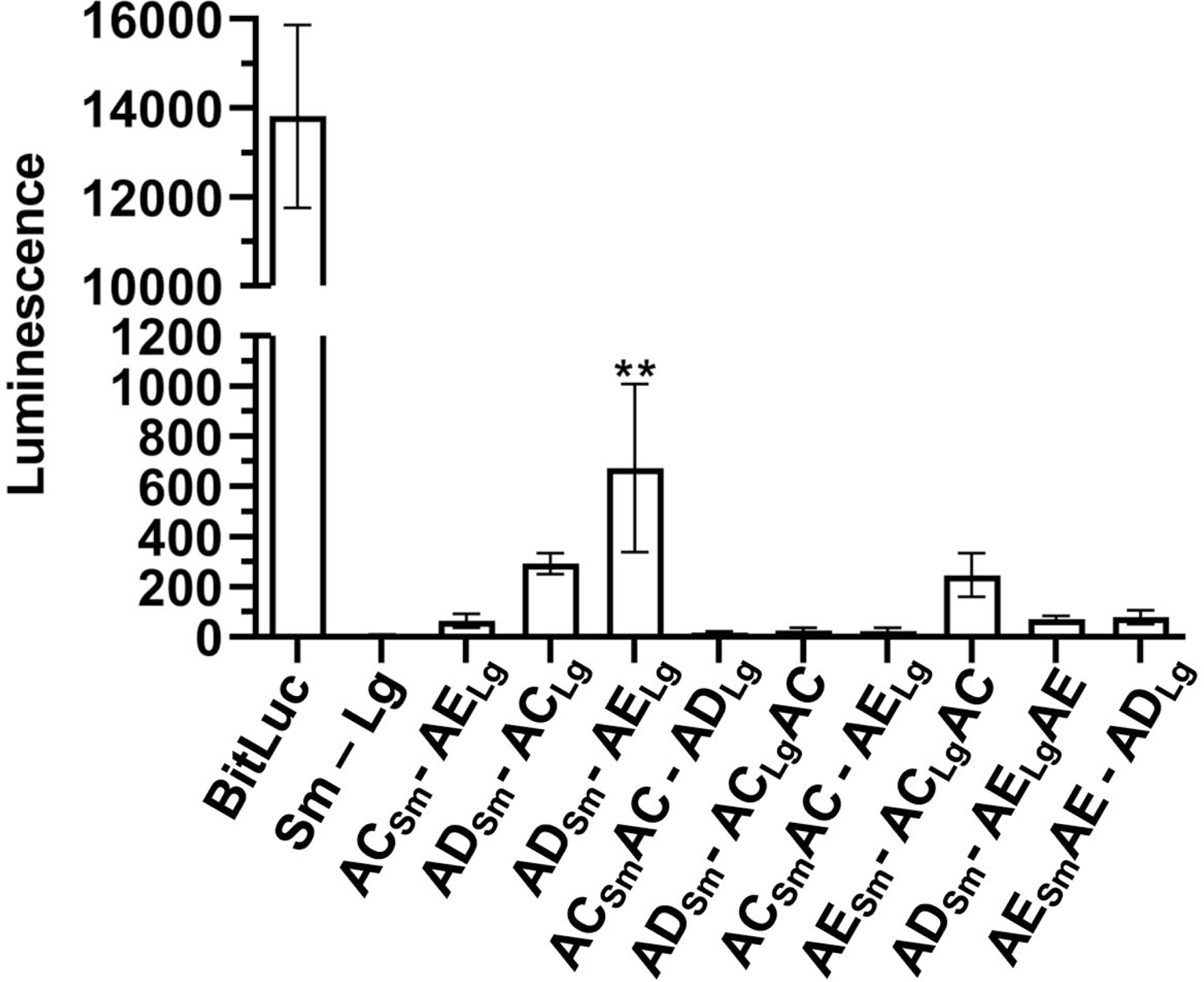
SpoVAD and SpoVAE interact in sporulating cultures *C. difficile* cells transformed with the indicated plasmids were grown on sporulation medium. Luminescence of the sporulating culture was assayed after 2 days of growth. Averages of 4 biological replicates are shown. ** p < 0.001 as determined by one-way ANOVA using Dunnett’s multiple comparisons test.

Because our data indicate that SpoVAD and SpoVAE interact during sporulation, it is plausible that these proteins would remain in close proximity in the metabolically dormant spore. We therefore tested the same protein interaction pairs in fully formed dormant spores. The strains were sporulated on sporulation medium, and spores purified as previously described.

Next, 1×10^8^ spores of each strain were assayed for luciferase activity (Figure 5). Similar to what we observed for a sporulating culture, the SpoVAD-SpoVAE interacting pair showed the highest level of interaction, while SpoVAC-SpoVAD, SpoVAD-SpoVAC, and SpoVAE-SpoVAClgAC showed a small but non-significant signal. Taken together, our results suggest that SpoVAD-SpoVAE interact when expressed in vegetative cell or in sporulating / dormant spores, while the other SpoVA proteins may weakly and / or transiently interact.

**Figure 5.**
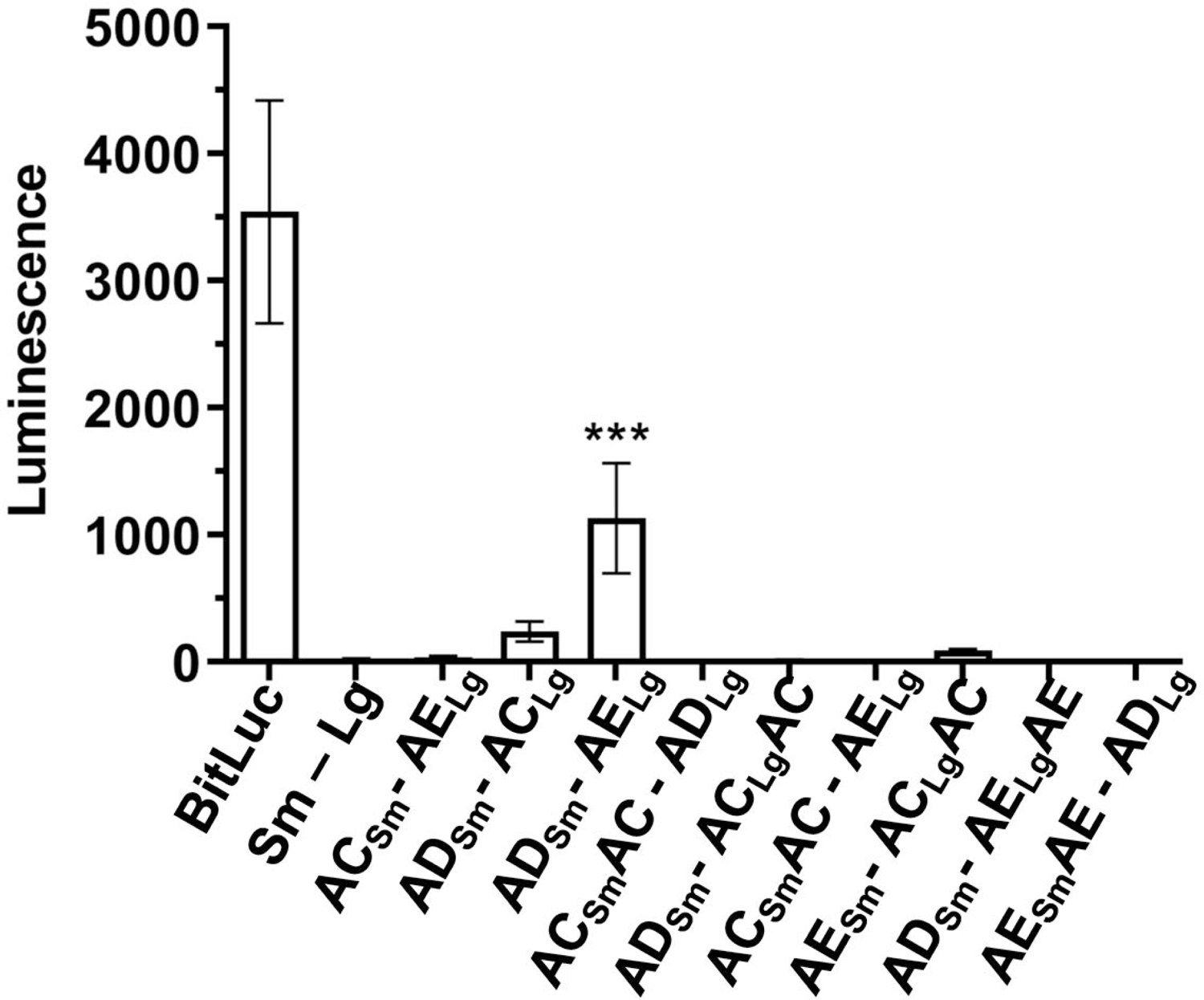
SpoVAD and SpoVAE demonstrate interact in dormant spores *C. difficile* cells transformed with the indicated plasmids were grown on sporulation medium and the resulting spores were purified and assayed for luminescence. Averages of 3 biological triplicates shown. *** p < 0.0001 as determined by one-way ANOVA using Dunnett’s multiple comparisons test.

We hypothesized that the SpoVA proteins that interact during DPA packaging would still interact upon spore germination. To test this, we exposed 1×10^8^ spores derived from the indicated strains to the germinants taurocholate and glycine in buffer and measured the luminescence signal over the period of 30 minutes. Similar to our observation in the sporulating culture and dormant spores, the SpoVAD-SpoVAE seems to interact the most strongly and gives the highest luminescence signal (Figure 6). The other interacting pairs, the C-terminal fusions and mid-protein fusions, also had an increase in signal. However, these signals did not reach statistical significance. But this and the similar signal increase in the previous assays using sporulating cultures and dormant spores suggests either a weak or transient interaction. We therefore conclude that SpoVAD and SpoVAE interact during all stages of *C. difficile* development cycle; in vegetative cells, sporulating cells, spores, and germinating spores.

**Figure 6.**
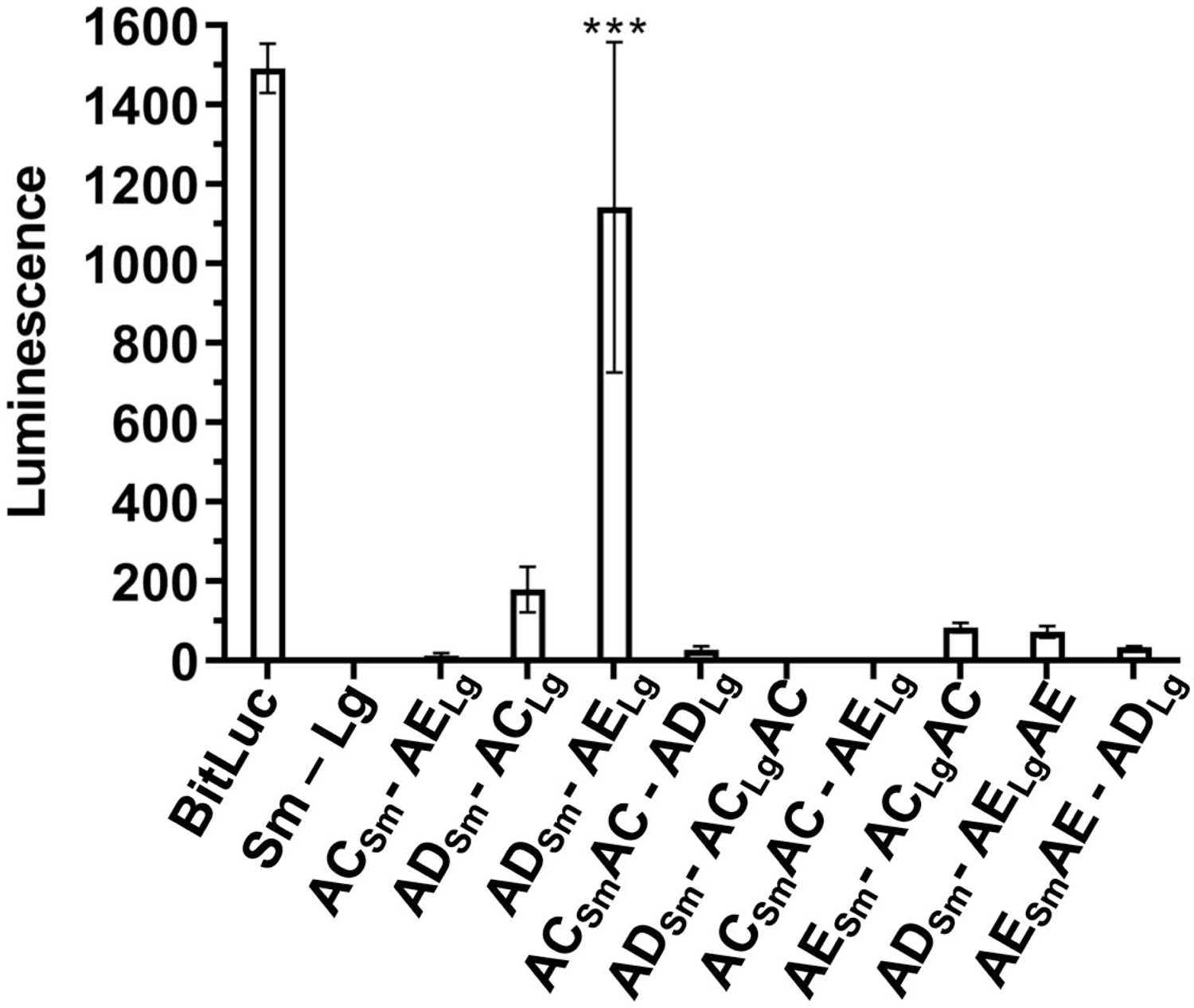
SpoVAD and SpoVAE interact during spore germination Spores derived from the strains containing the indicated plasmids were germinated in the presence of TA and glycine. Luminescence was measured over the period of 30 minutes, but only T = 15 minutes shown for clarity. Averages of 3 biological triplicates shown. *** p < 0.0001 as determined by one-way ANOVA using Dunnett’s multiple comparisons test.

## Discussion

*C. difficile* spores, like most spores from endospore-forming organisms, are notable for their resilience to heat. Peak natural environment temperatures and even temperatures that are commonly recommended for heat treatment of foodstuffs are often insufficient to destroy the spores or render them incapable of germination (42). This heat resistance, along with other characteristics of spores that provide resistance to UV radiation, desiccation, and resistance to common disinfectants, makes *C. difficile* (or other spore-forming organisms) difficult to eradicate (43). The spore resistance to heat is largely the consequence of large amounts of DPA in the spore core that replaces the majority of water and comprises 5-15% of dry weight of the spore (44). Since spores are metabolically dormant, in order for metabolic processes to initiate, this DPA must be released from the spore core in exchange for water during the initial stages of spore germination.

The initiation of germination is a non-reversible process and the release of DPA in exchange for water is tightly regulated by a mechanism. To initiate *C. difficile* spore germination, the Csp-type germinant receptors, composed of a hypothesized CspB, CspA, and CspC protein complex, become activated by bile acid and amino acid germinants. Germinant activation of the germinant receptors leads to the processing of the inhibitory pro-peptide from the cortex lytic enzyme SleC (36, 45). Activated SleC degrades the spore cortex layer resulting in the activation of the SpoVAC mechanosensing protein and DPA release from the core (12, 27). Though, the importance of *spoVAC* is known in both *B. subtilis* and *C. difficile*, the roles of *spoVAD* and *spoVAE* have not been tested in *C. difficile*. Here, we find that the mutations in *C. difficile spoVAD* and *spoVAE* prevent DPA packaging into the spore core. This is similar to prior observations with *C. difficile spoVAC* mutants (27, 32).

Because a mutation in any member of the *C. difficile spoVA* operon resulted in a similar phenotype, we hypothesized that the products of this operon form a complex and interact during DPA packaging. Because SpoVAC is a transmembrane protein that is embedded in the inner spore membrane, and SpoVAE is predicted to have transmembrane domains, this complex should be located at, or near, the inner spore membrane. Even though SpoVAD is not predicted to have a transmembrane domain, consistent with its hypothesized role as a DPA binding protein (29), it should be located near the inner spore membrane during DPA packaging in order to transfer DPA via SpoVAC into the spore core. Using the recently developed split luciferase system for studying protein-protein interactions in *C. difficile* (39), we detected a large increase in luminescence for SpoVAD and SpoVAE and smaller, but non-significant increases in other interaction pairs, in vegetative cells, sporulating cells, dormant spores, and germinating spores. We therefore propose a model in which the DPA, synthesized in the mother cell in the late stage of sporulation, is packaged into the spore by the interaction between all 3 SpoVA proteins. In this model, SpoVAD acts as a DPA binding protein, SpoVAC as a channel through which DPA passes into the spore core, and SpoVAE acting as an accessory protein (Figure 7). Consistent with the predictions arising from this model, we first observed interaction between all 3 protein pairs in vegetative cells (i.e., SpoVAC-SpoVAE, SpoVAD-SpoVAC, and SpoVAD-SpoVAE). In our assay, the protein pair expression was driven by an aTc-induced pTet promoter which likely induces protein expression above the level of baseline expression in vegetative *C. difficile* cells, but this served to show that the interaction partners could interact, and to discover any interactions that may be transient, temporary, or at low incidence rate in the uninduced culture. We also discovered such interactions in sporulating cultures, in the dormant spores, and in the germinating spores. We found that in all 3 stages of spore development the strongest interacting partners were SpoVAD and SpoVAE. This has given us confidence in our model because SpoVAE has predicted transmembrane domains and is found in the inner spore membrane of *B. subtilis* spores (29, 30). Though SpoVAD has no predicted transmembrane domains, our data suggests it is situated on the inner spore membrane with SpoVAE.

**Figure 7.**
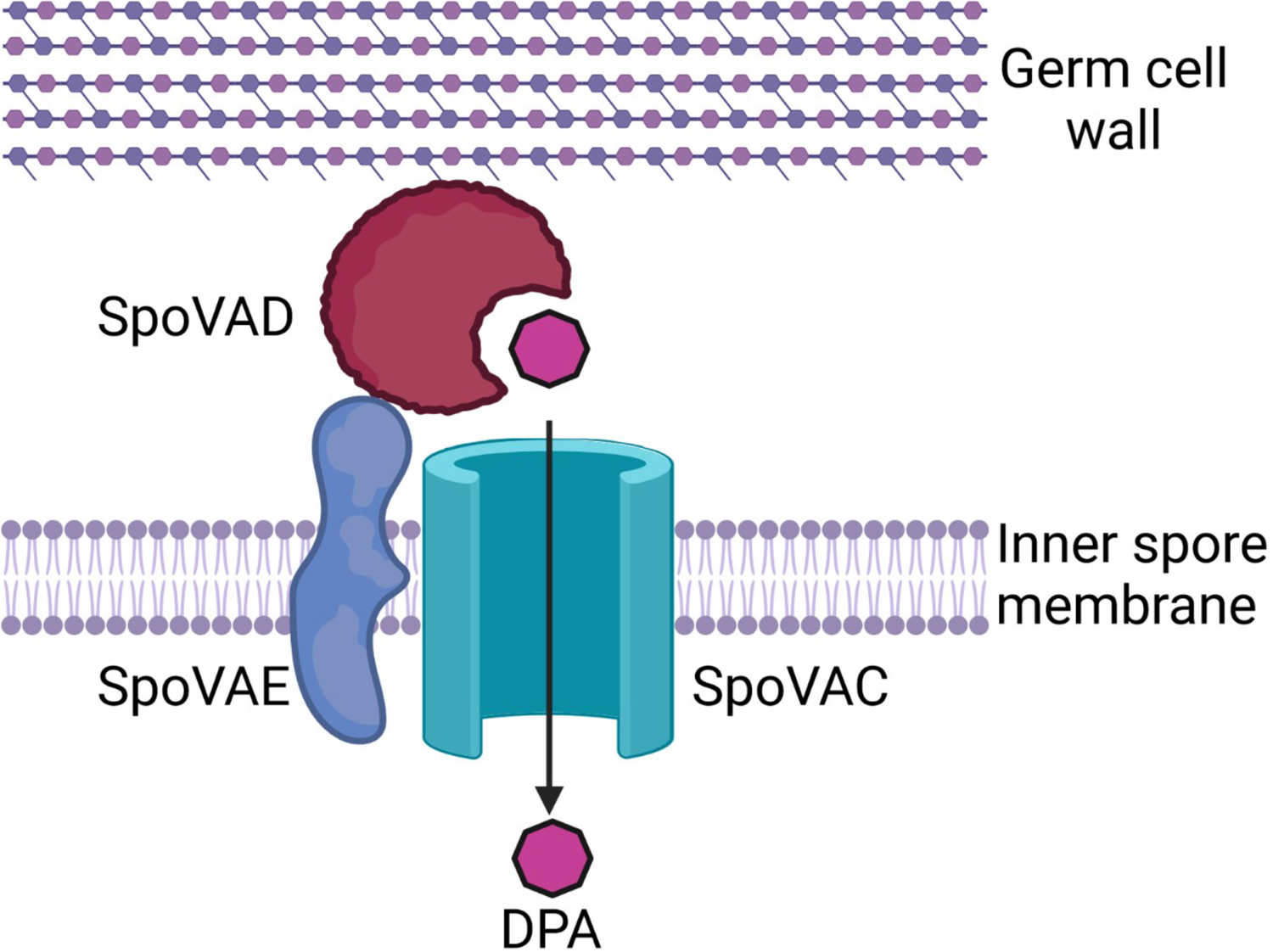
Proposed model SpoVA protein interaction at the inner spore membrane by which DPA produced by the mother cell is packaged into the spore during sporulation and released in the initial stages of germination. Created with BioRender.com

Because the expression of the fusions was driven by an inducible promoter, the data likely represent the signal at the maximum expression levels, which may be why the SpoVAC-SpoVAD interaction signal is quite high. Because the mother cell produces large amounts of DPA during the late stages of sporulation, the trafficking of DPA by SpoVAD to SpoVAC, and their interaction, must be a high-incidence event, if perhaps temporary. After sporulation is completed, the assays showed the highest interaction signal for SpoVAE-SpoVAD, suggesting that this pair forms a more stable interaction.

In the model spore-forming organism *B. subtilis*, the *spoVA* operon is composed of 8 genes, while the *C. difficile spoVA* operon is considerably less complex and encodes only 3 orthologues, making it a convenient system for study. Since our data suggests that all of the *C. difficile* SpoVA proteins may interact, we hypothesize that the protein products of a more complex *B. subtilis spoVA* operon form a complex, as well.

## Materials and methods

### Bacteria and strains

*C. difficile* strains were grown at 37 °C in an anaerobic chamber (Coy Laboratories, model B, >4% H_2_, 5% CO_2_, 85% N_2_), on brain heart infusion agar supplemented by 5 g / liter yeast extract and 0.1% L-cysteine (BHIS). *E. coli* DH5α (46) was grown on LB medium. Chloramphenicol (20 µg / mL), thiamphenicol (10 µg / mL), kanamycin (50 µg / mL), ampicillin (100 μg / mL) were added where indicated. Deletion mutants were selected on *C*. *difficile* minimal medium (CDMM) supplemented with 5 μg / mL 5-fluoroorotic acid (FOA) and 20 μg / mL uracil.

### Construction of *spoVAD* and *spoVAE* mutants

Deletion mutations were introduced using the established *pyrE*-mediated allelic exchange technique (33). Briefly, 1kb upstream and downstream DNA regions that surround *spoVAD* and *spoVAE* (including the first 30 bp of the 5’ and 3’ of the gene) were amplified using primers spoVAD_ndeI_L, spoVAD_ndeI_R, spoVAD_KO_RHA_For, spoVAD_xhoI_R for *spoVAD* and spoVAE_ndeI_L, 250_Downstream_VAE, 250_Upstream_VAE, spoVAE_xhoI_R for *spoVAE*, assembled using PCR stitching, inserted by Gibson assembly (47) into pMTL-YN4 plasmid digested with NdeI and XhoI, yielding plasmids pMB02 and pMB04. The plasmids were then transformed into *E. coli* DH5α. These plasmids were subsequently transformed into *E. coli* HB101 pRK24 and grown on LB medium supplemented with chloramphenicol and ampicillin.

The resulting strain was grown overnight and then mixed with the *C. difficile* CRG2359 strain grown in BHIS in anaerobic chamber. The conjugation mixtures were spotted onto BHI plates and allowed to grow for 24 hours. Subsequently, the cells were washed with BHIS and the slurry was transferred onto BHIS medium supplemented with thiamphenicol (for plasmid maintenance) and kanamycin (to counter-select *E. coli* growth) [BHIS(TK)]. Individual colonies were passaged several times onto BHIS(TK) supplemented with uracil [BHIS(TKU)] to encourage the single crossover events. Growth was then transferred to CDMM medium supplemented with FOA and uracil to select for colonies that have excised the plasmid from the chromosome.

Thiamphenicol-sensitive colonies were tested for desired mutation by PCR and confirmed by sequencing. The wild-type *pyrE* allele was restored using the same technique. The resulting *C. difficile* Δ*spoVAD* strain was renamed MB03 and the *C. difficile* Δ*spoVAE* strain MB04. The mutations were complemented by introduction of a wild-type gene under the control of a native promoter on a plasmid.

### Construction of split luciferase interaction plasmids

Plasmids for luciferase assays were purchased from Addgene (ID: 105494 – 105497) and plasmids for testing interaction in vegetative cells were constructed following previously established protocols (39, 48). For these plasmids, *spoVAC, spoVAD,* and *spoVAE* was amplified using *C. difficile* R20291 DNA as a template to create fragments with the appropriate overlap for SmBit reporter fragment fusions (5’ SpoVAC SmBit Gibson / 3’ SpoVAC SmBit Gibson for *spoVAC,* 5’ SpoVAD SmBit Gibson / 3’ SpoVAD SmBit Gibson for *spoVAD,* 5’ SpoVAE SmBit Gibson / 3’ SpoVAE SmBit Gibson for *spoVAE*). These fragments were then introduced by Gibson assembly into plasmid pAP118 digested with SacI / XhoI. Using the same approach, fragments for LgBit reporter fusions were created (5’ SpoVAC LgBit Gibson / 3’ SpoVAC LgBit Gibson, 5’ SpoVAD LgBit Gibson / 3’ SpoVAD LgBit Gibson, 5’ SpoVAE LgBit Gibson / 3’ SpoVAE LgBit Gibson). These fragments were then ligated into the above plasmids, digested with PvuI / NotI, yielding the interaction plasmids. To construct the negative control interaction plasmids, plasmid pAF256 was digested with SacI / XhoI, plasmid pAF257 digested with PvuI / NotI, and the required single fusion fragments were introduced into them by Gibson assembly. To construct the negative control interaction plasmid, with only SmBit and LgBit reporters without the protein fusion, LgBit reporter fragment was amplified from pAF256 using primers 5’ LgBit Gibson / 3’ LgBit Gibson and inserted by Gibson assembly (47) into plasmid pAF257 digested with PvuI / BamHI. This yielded plasmids pMB45 through pMB62 with all combinations of interacting partners and negative controls needed for vegetative cell expression. Next, we created plasmids with a native *spoVAC* promoter to drive the expression of the requisite constructs from the native promoter. For SpoVAC-SpoVAE we amplified the promotor region fragment from a wild-type template using primers 5’ SpoVAC prm / 3’ SpoVAC prm-SpoVAC, and the fusion partner sequence from pMB45 using primers 5’ SpoVAC RBS / 3’ LgBit-pJS116. For SpoVAD-SpoVAC, we amplified the promotor region from a wild-type template using primers 5’ SpoVAC prm / 3’ spoVAC pr. - spoVAD Gibson, and the fusion partner sequence from pMB46 using primers 5’ SpoVAC prm-SpoVAD / 3’ LgBit-pJS116. For SpoVAD-SpoVAE we amplified the promoter region from a wild-type template using primers 5’ SpoVAC prm / spoVAC pr. - spoVAD Gibson and the fusion partner sequence from pMB48 using primers 5’ SpoVAC prm-SpoVAD / 3’ LgBit-pJS116. Fragments were introduced by Gibson assembly into pMTL84151 digested with NotI / HinDIII. This yielded plasmids pMB72 (SpoVAC-SpoVAE), pMB73 (SpoVAD-SpoVAC), and pMB75 (SpoVAD-SpoVAE). We also created the plasmids where the SmBit or LgBit reporter fragment is inserted into an extra cytoplasmic-facing segment of SpoVAC or SpoVAE. For AC_Sm_AC-AD_Lg_ we amplified fragments from wild-type template using primers 5’ SpoVAC downstream (linker-SmBit) / 3’ SpoVAC downstream (linker-SmBit) and primers 5’ SpoVAC prm / 3’ SpoVAC upstream (linker-SmBit), from pMB54 template using primers 5’ SpoVAC-SpoVAD / 3’ LgBit-pJS116, and from pMB51 template using primers 5’ linker-SmBit (SpoVAC) / 3’ linker-SmBit (SpoVAC). For AD_Lg_-AC_Sm_AC we amplified fragments from the wild-type template using primers 5’ SpoVAC prm / 3’ spoVAC pr. - spoVAD Gibson, primers 5’ SpoVAC LgBit Gibson / 3’ SpoVAC upstream (linker-SmBit), primers 5’ linker-LgBit (SpoVAC) / 3’ Linker-LgBit (SpoVAC), from pMB51 template using primers 5’ SpoVAC prm-SpoVAD / 3’ SmBit-SpoVAC RBS, and from pMB54 template using primers 5’ linker-SmBit (SpoVAC) / 3’ Linker-LgBit. For AC_Sm_AC-AE_Lg_ we amplified fragments from the wild-type template using primers 5’ SpoVAC prm / 3’ SpoVAC upstream (linker-SmBit), primers 5’ SpoVAC downstream (linker-SmBit) / 3’ SpoVAC downstream (linker-SmBit) – SpoVAE, from pMB51 template using primers 5’ linker-SmBit (SpoVAC) / 3’ linker-SmBit (SpoVAC), and from pMB54 template using primers 5’ SpoVAC-SpoVAE / 3’ Linker-LgBit. For AE_Sm_-AC_Lg_AC we amplified fragments from the wild-type template using primers 5’ SpoVAC prm / 3’ spoVAC pr. - spoVAE Gibson, primers 5’ SpoVAC LgBit Gibson / 3’ SpoVAC upstream (linker-SmBit), primers 5’ linker-LgBit (SpoVAC) / 3’ Linker-LgBit (SpoVAC), from pMB52 template using primers 5’ spoVAC pr. - spoVAE Gibson / 3’ SmBit-SpoVAC RBS, and from pMB54 template using primers 5’ linker-SmBit (SpoVAC) / 3’ Linker-LgBit (SpoVAC). For AD_Sm_-AE_Lg_AE we amplified fragments from the wild-type template using primers 5’ SpoVAC prm / 3’ spoVAC pr. - spoVAD Gibson, primers 5’ SpoVAE LgBit Gibson / 3’ spoVAE-SpoVADSm, primers 5’ SpoVAE-plasmid LgBit Mid / 3’ SpoVAE-plasmid LgBit Mid, from pMB75 template using primers 5’ SpoVAC prm-SpoVAD / 3’ spoVAD-SmBit, and from pAP118 template using primers 5’ SpoVAE LgBit Mid / 3’SpoVAE LgBit Mid. For AE_S / m_AE-AD_Lg_ we amplified fragments from the wild-type template using primers 5’ SpoVAC prm / 3’ spoVAC pr. - spoVAE Gibson, primers 5’ spoVAC pr. - spoVAE Gibson / 3’ SpoVAESm-SpoVADLg, primers 5’ SpoVAE Smbit Mid / 3’ SpoVAE Smbit Mid, from pMB50 template using primers 5’ SpoVAE-SmBit / 3’ SpoVAE-SmBit, and from pMB56 template using primers 5’ SpoVAE SmBit mid - SpoVAD LgBit / 3’ LgBit-pJS116. The fragments were introduced by Gibson assembly (47) into the pMTL84151 plasmid digested with NotI / HinDII, yielding plasmids pMB82 – pMB85, and pMB88 – pMB89. All strains and plasmids in this study are listed in Table 1. Primers used to construct the strains and plasmids are listed in Table 2.

**Table 1.** List of strains and plasmids used in this study

| ***C. difficile* strains** | **Description** | **Reference** |
| --- | --- | --- |
| R20291 | Wild type, ribotype 027 | (50) |
| CRG2359 | R20291 $\Delta pyrE$ | (33) |
| RS19 | CRG2359 restored *pyrE* | (36) |
| MB03 | $\Delta spoVAD$ strain | This study |
| MB04 | $\Delta spoVAE$ strain | This study |
| **Other strains** |  |  |
| *E. coli* DH5α | Cloning strain | (46) |
| *E. coli* HB101 pRK24 | Conjugal donor strain for *C. difficile* | (51) |
| **Plasmid** | **Description** | **Reference** |
| pMTL-YN4 | Backbone used to make deletions in R20291 by allelic exchange | (33) |
| pMTL84151 | Backbone used to make complementing plasmids and spore luciferase interaction plasmids | (52) |
| pMB02 | To create $\Delta spoVAD$ deletion | This study |
| pMB04 | To create $\Delta spoVAE$ deletion | This study |
| pMB34 | *spoVAD* complementing plasmid | This study |
| pMB35 | *spoVAE* complementing plasmid | This study |
| pAP118 | Plasmid encoding an aTC inducible HupA-SmBiT and HupA-LgBiT fusions | (39) |
| pAF256 | Plasmid encoding an aTC inducible HupA-SmBiT and LgBiT | (39) |
| pAF257 | Plasmid encoding an aTC inducible SmBiT and HupA-LgBiT | (39) |
| pAF259 | Plasmid encoding an aTC inducible BitlucOpt | (39) |
| pMB50 | Luciferase *spoVAC* / Smbit-LgBiT | This study |
| pMB51 | Luciferase *spoVAD* / Smbit-LgBiT | This study |
| pMB52 | Luciferase *spoVAE* / Smbit-LgBiT | This study |
| pMB53 | Luciferase Smbit-*spoVAC* / LgBiT | This study |
| pMB54 | Luciferase Smbit-*spoVAD* / LgBiT | This study |
| pMB55 | Luciferase Smbit-*spoVAE* / LgBiT | This study |
| pMB62 | Luciferase *spoVAC* / Smbit-SpoVAC / LgBiT | This study |
| pMB45 | Luciferase *spoVAC* / Smbit-SpoVAE / LgBiT | This study |
| pMB46 | Luciferase *spoVAD* / Smbit-SpoVAC / LgBiT | This study |
| pMB47 | Luciferase spoVAD / Smbit-SpoVAD / LgBit | This study |
| pMB48 | Luciferase spoVAD / Smbit-SpoVAE / LgBit | This study |
| pMB49 | Luciferase spoVAE / Smbit-SpoVAC / LgBit | This study |
| pMB57 | Luciferase spoVAE / Smbit-SpoVAE / LgBit | This study |
| pMB60 | Luciferase spoVAC / Smbit-SpoVAD / LgBit | This study |
| pMB61 | Luciferase Smbit-LgBit | This study |
| pMB72 | spoVAC-SmBit / spoVAE-LgBit<br>luciferase spore complementation | This study |
| pMB73 | spoVAD-SmBit / spoVAC-LgBit<br>luciferase spore complementation | This study |
| pMB75 | spoVAD-SmBit / spoVAE-LgBit<br>luciferase spore complementation | This study |
| pMB79 | HupA-SmBit / HupA-LgBit luciferase<br>spore complementation | This study |
| pMB81 | BitLuc luciferase spore<br>complementation | This study |
| pMB82 | SpoVAC-SmBit-SpoVAC / SpoVAD-LgBit; small luciferase reporter<br>inserted mid-AC gene, ADLg partner | This study |
| pMB83 | SpoVAD-SmBit / SpoVAC-LgBit-SpoVAC; large luciferase reporter<br>inserted mid-AC gene, ADSm partner | This study |
| pMB84 | SpoVAC-SmBit-SpoVAC / SpoVAE-LgBit; small luciferase reporter<br>inserted mid-AC gene, AELg partner | This study |
| pMB85 | SpoVE-SmBit / SpoVAC-LgBit-SpoVAC; large luciferase reporter<br>inserted mid-AC gene, AESm partner | This study |
| pMB87 | Sm-Lg luciferase spore interaction<br>plasmid, negative control. | This study |
| pMB88 | SpoVAD-Sm / SpoVAE-Lg-SpoVAE;<br>large luciferase reporter inserted mid-AE gene, ADSm partner | This study |
| pMB89 | SpoVAE-Sm-SpoVAE / SpoVAD-Lg;<br>small luciferase reporter inserted<br>mid-AE gene, ADLg partner | This study |

**Table 2.**
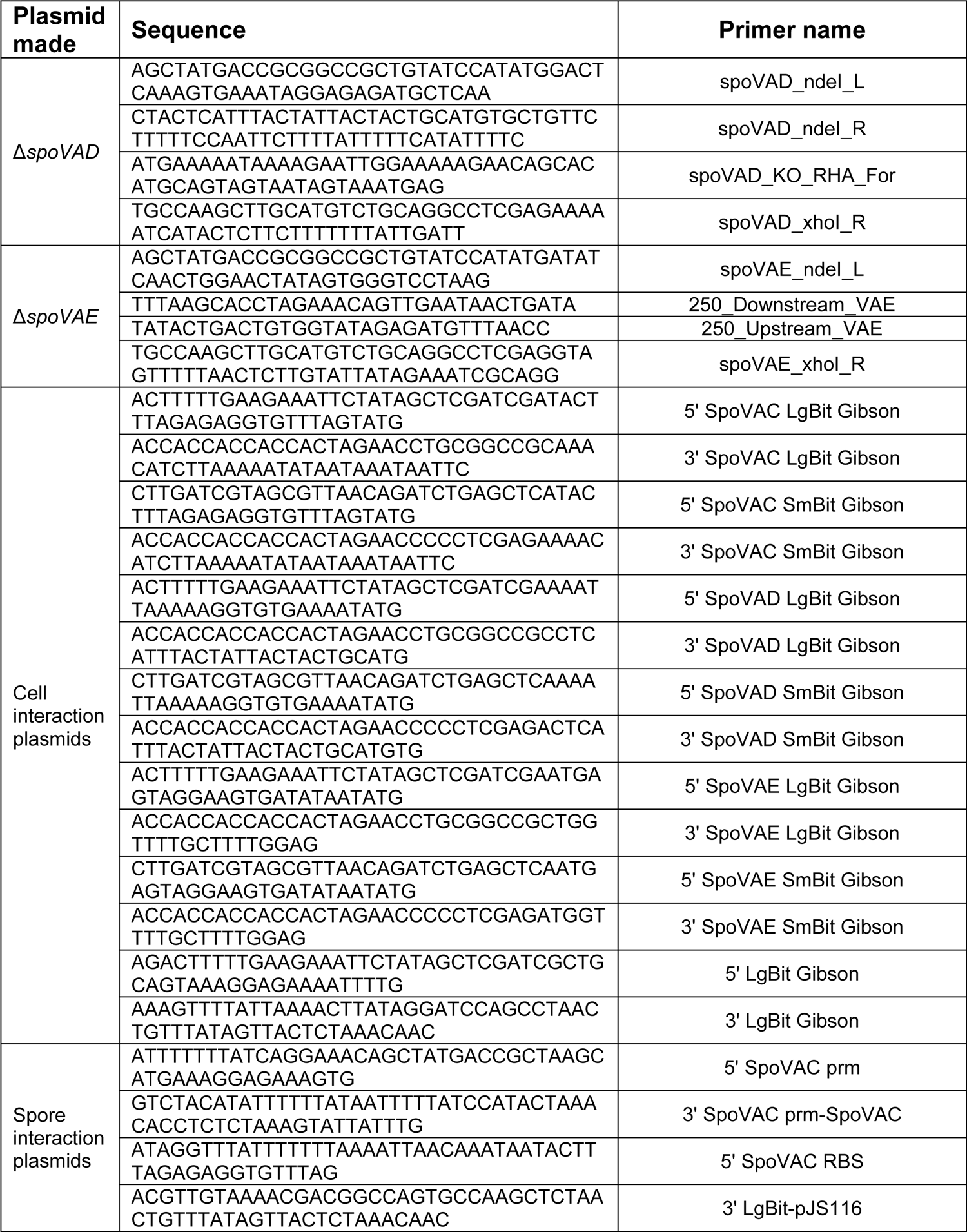
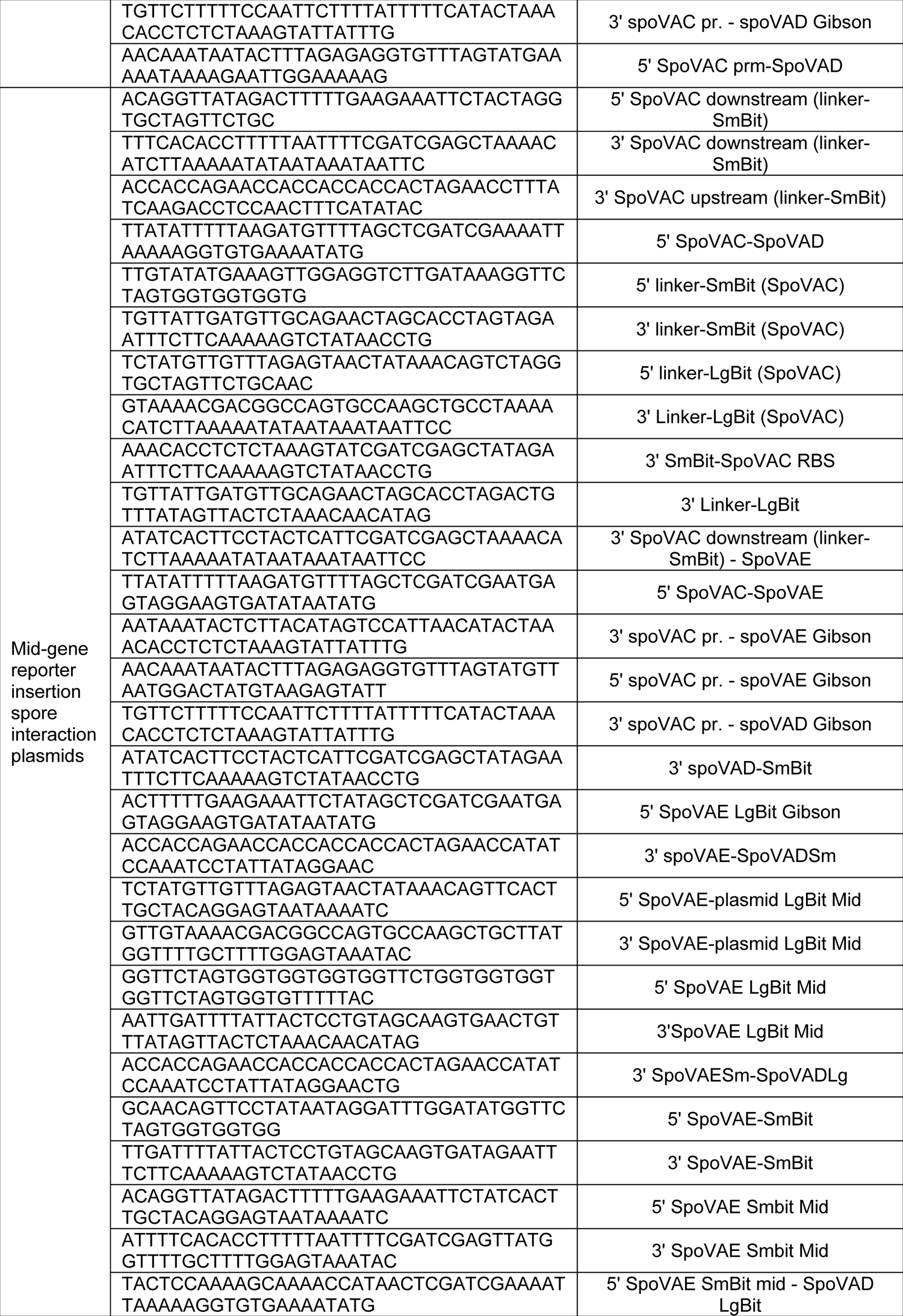
List of oligonucleotides used in this study

### Spore purification

Spores were purified as previously described (12, 23, 27, 34). Briefly, strains were grown on 70:30 sporulation medium. After 5 days, growth from 2 plates each was scraped into 1 mL dH_2_O in microcentrifuge tubes and left overnight at 4 °C. The cultures were then resuspended in the dH_2_0 in the same microcentrifuge tubes, centrifuged at >14,000 x g for 10 minutes, the top layer containing vegetative cells and cell debris was removed by pipetting, and the rest of the sediment resuspended in fresh dH_2_0. The tubes, again, were centrifuged for 1 minute at >14,000 x g, the top layer removed, and the sediment resuspended. This was repeated 5 more times, combining the sediment from 2 tubes into one. The spores were then separated from the cell debris by centrifugation through a 50% sucrose gradient for 20 minutes at 4 °C and 3,500 x g. The resulting spore pellet was then washed 5 times with dH_2_0, resuspended in 1 mL dH_2_0, and stored at 4 °C until use.

### DPA and OD germination assays

Ca-DPA release was measured using a SpectraMax M3 plate reader for 1 hr at 37 °C with excitation at 270 nM and emission at 545 nM and with a 420 nM cutoff. Spores were heat activated at 65 °C for 30 minutes and suspended in water at an OD_600_ = 50. The spores were then added to final OD of 0.5 in 100 μL final volume of HEPES buffer pH 7.5, containing 100 mM NaCl, 10 mM TA, 30 mM glycine, and 250 μM Tb^3+^ in a 96 well plate. OD_600_ was monitored using the same plate reader at 37 °C for 1 hr. The heat-activated spores were added to a final OD of 0.5 in in the same HEPES buffer composition as in Ca-DPA release assay, omitting Tb^3+^ (36, 49).

### Luciferase assays

Assays involving vegetative cells were performed using the previously published protocols (39). Briefly, strains were grown overnight in liquid BHIS supplemented with thiamphenicol. The following day the liquid cultures were diluted to an OD_600_ = 0.05 and allowed to grow to an OD_600_ = 0.3 – 0.4. Next, 100 μL was removed and placed into wells of a transparent 96 well plate, suitable for OD_600_ and another 100 μL placed into white flat-bottom 96-well plate for the luminescence assay. The rest of the cultures were induced with 200 ng / mL of aTc for 1 hour. 20 μL of NanoGlo luciferase (Promega N1110) was added to each sample well to measure luciferase activity using SpectraMax M3 plate reader in the luminescence mode, using all channels, and 0.1 second sampling time. After 1 hour, the induced cultures were assayed for OD and luciferase activity in the same way. The luciferase activity was normalized to culture optical density at OD_600_.

### Western blot

Solutions of 1×10^8^ spores of *C. difficile* R20291, *C. difficile* Δ*spoVAD,* and *C. difficile* Δ*spoVAE* were prepared and 100 µL of each was incubated in HEPES buffer pH 7.5, containing 100 mM NaCl, 10 mM TA, 30 mM glycine for 15 minutes to induce germination. These samples and equal amounts of non-germinated control spore solutions were boiled for 20 minutes in 2x NuPage buffer at 95 °C. Then, 10 μL of each sample was separated on 10% SDS-PAGE gel. The protein was transferred to PVDF membrane and then blocked overnight at 4 °C with 5% milk powder dissolved in TBST. The membrane was washed thrice for 20 minutes at room temperature with TBST and then probed for 1 hour in 5% milk dissolved in TBST with anti-SleC antibodies at room temperature. The membrane was then washed thrice for 20 minutes at room temperature with TBST before labeling with anti-rabbit IgG secondary antibody. The membrane was again washed and then incubated for 5 minutes with Pierce ECL Western Blotting Substrate (ThermoScientific), overlaid with X-ray film, exposed and developed.

### Statistical analysis

Data represents results from at least 3 independent and the error bars represent standard errors of the means. One-way ANOVA followed by Tukey’s or Dunnett’s multiple comparisons test, as indicated, was performed using GraphPad Prism version 9.0.2 (161) for Windows, (GraphPad Software, San Diego, California USA).

## Acknowledgments

This project was supported by awards 5R01AI116895 and 1U01AI124290 to J.A.S. from the National Institute of Allergy and Infectious Diseases. The content is solely the responsibility of the authors and does not necessarily represent the official views of the NIAID. The funders had no role in study design, data collection and interpretation, or the decision to submit the work for publication.

